# Endothelial Unc5B controls blood-brain barrier integrity

**DOI:** 10.1101/2021.05.06.442974

**Authors:** Kevin Boyé, Luiz Henrique Geraldo, Jessica Furtado, Laurence Pibouin-Fragner, Mathilde Poulet, Doyeun Kim, Bryce Nelson, Yunling Xu, Laurent Jacob, Nawal Maissa, Bertrand Tavitian, Dritan Agalliu, Lena Claesson-Welsh, Susan L Ackerman, Anne Eichmann

**Affiliations:** Cardiovascular Research Center, Department of Internal Medicine, Yale University School of Medicine, New Haven CT, USA; Paris Cardiovascular Research Center, Inserm U970, Université Paris, France; Medicinal Bioconvergence Research Center, Yonsei University, Incheon, Republic of Korea; Department of Pharmacology, Cancer Biology Institute, Yale University School of Medicine, New Haven, CT, USA; Departments of Neurology and Pathology and Cell Biology, Columbia University Irving Medical Center, New York NY, USA; Department of Immunology, Genetics and Pathology, Uppsala University, Uppsala, Sweden; Division of Biological Sciences Section of Neurobiology and Department of Cellular and Molecular Medicine, University of California San Diego and Howard Hughes Medical Institute, La Jolla CA, USA; Department of Molecular and Cellular Physiology, Yale University School of Medicine, New Haven CT, USA

## Abstract

Blood-brain barrier (BBB) integrity is critical for proper function of the central nervous system (CNS). Here, we showed that the endothelial Netrin1 receptor Unc5B controls BBB integrity by maintaining Wnt/β–catenin signaling. Inducible endothelial-specific deletion of Unc5B in adult mice led to region and size-selective BBB opening. Loss of Unc5B decreased BBB Wnt/β–catenin signaling, and β–catenin overexpression rescued *Unc5B* mutant BBB defects. Mechanistically, Netrin1 enhanced Unc5B interaction with the Wnt co-receptor LRP6, induced its phosphorylation and activated Wnt/β–catenin downstream signaling. Intravenous delivery of antibodies blocking Netrin1 binding to Unc5B caused a transient disruption of Wnt signaling and BBB breakdown, followed by neurovascular barrier resealing. These data identify Netrin-Unc5B signaling as a novel regulator of BBB integrity with potential therapeutic utility for CNS diseases.

## Introduction

The BBB protects the brain from toxins and pathogens and maintains homeostasis and proper function of the CNS^1,2^. Wnt/β-catenin signaling maintains BBB integrity via expression of either Wnt7a,7b or Norrin ligands, which bind to multiprotein receptor complexes including Frizzled4 and LRP6 on brain endothelial cells (ECs) in distinct CNS regions ^3–5^. Receptor activation causes β–catenin stabilization and nuclear translocation to induce expression of a BBB-specific gene transcription repertoire, including the tight junction (TJ)-associated protein Claudin5 that suppresses paracellular permeability, while inhibiting expression of the permeability protein PLVAP that forms the diaphragm in EC fenestrae and transcytotic vesicles^6–9^. Whether Wnt/β-catenin signaling could be modulated to open the BBB “on-demand” or to restore its integrity when damaged, is unknown.

Here, we have identified the endothelial Unc5B receptor and its ligand Netrin1 as novel regulators of Wnt/β–catenin signaling at the BBB. Unc5B is a transmembrane receptor for Netrin1^10,11^, Robo4^12,13^ and Flrt2^14,15^ that is predominantly expressed in ECs in mice and humans^16^. Global *Unc5B* knockout in mice is embryonically lethal due to vascular defects^14,16^, demonstrating that Unc5B has important functions in vascular development. Whether Unc5B signaling is required in postnatal mice or in adults remained unknown.

## Results

### Unc5B controls BBB development and maintenance

We generated tamoxifen (TAM)-inducible, endothelial-specific *Unc5B* knockout mice by crossing *Unc5B^fl/fl^* mice (**Supp. Fig. 1a**) with *Cdh5Cre^ERT2^* mice^17^ (hereafter Unc5BiECko), which deletes in ECs. Gene deletion was induced by TAM injection in neonates between postnatal day (P)0-P2, and qPCR revealed efficient *Unc5B* deletion (**Supp. Fig. 1b,c**). Interestingly, neonatal TAM injection induced seizures and lethality of Unc5BiECko mice around P15 (**Supp. Fig. 1d, Supp videos 1-4**), indicating an abnormal excitability of the neuronal network that may result from a BBB failure^2^. Intraperitoneal injection of a fluorescent tracer cadaverine (MW 950Da) into P5 mice and analysis of tracer leak 2hrs later revealed widespread tracer extravasation into the brain of P5 Unc5BiECko mice, which confirmed that Unc5B deletion impaired BBB development (**Supp. Fig. 1,e**).

To determine if Unc5B also controlled BBB integrity in adults, we induced gene deletion and probed BBB integrity 7 days later by intravenous (i.v.) injection of fluorescent tracers (**Fig.1a**). Cadaverine remained inside the vasculature of TAM injected Cre-littermate controls but leaked into the Unc5BiECko brain (**Fig. 1b**). Cadaverine leakage was observed in several regions of adult Unc5BiECko brains, including the retrosplenial and piriform cortex, hippocampus, hypothalamus, thalamus, striatum and cerebellum, while other cortical areas such as the posterior parietal association areas and the primary somatosensory cortex displayed an intact BBB (**Fig. 1c**). Injection of fluorescent dextrans of increasing molecular weights showed that both 10kDa and 40kDa dextrans had a higher permeability across the BBB in Unc5BiECko brains compared to controls, whereas 70kDa dextran did not cross the BBB, suggesting a size selective defect of BBB leakage for proteins greater than 40kDa in Unc5BiECko mice (**Fig. 1d**). The vascular permeability to cadaverine and 40kDa dextran in other Unc5BiECko organs was similar to controls (**Supp. Fig.1f**), demonstrating that Unc5B has a CNS-selective BBB protective function in adult mice, which may be due to its enriched expression in adult brain endothelium when compared to endothelium of other organs^18^.

**Figure 1:**
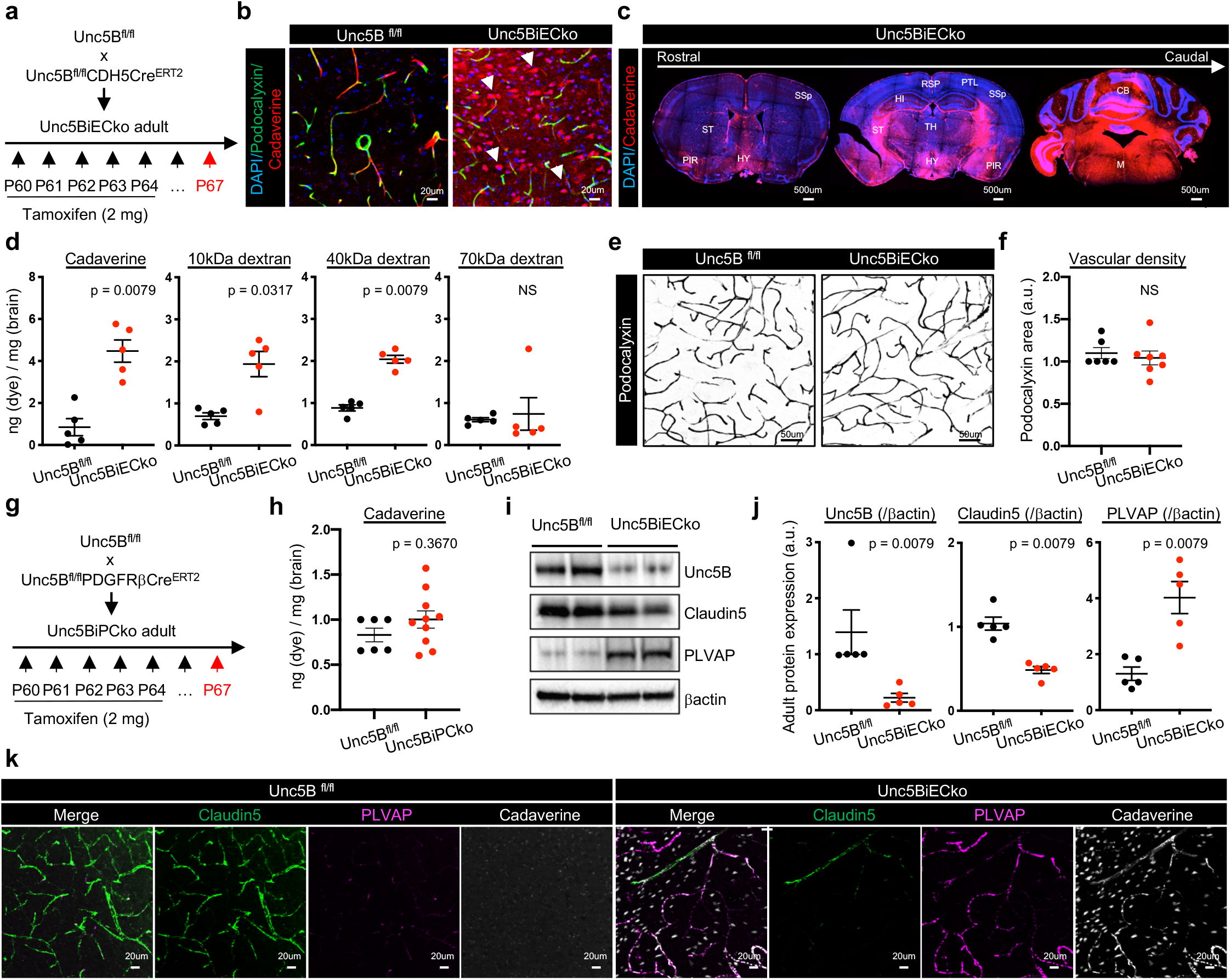
Unc5B controls BBB integrity. (a) *Unc5B* gene deletion strategy using tamoxifen (TAM) injection in adult mice. (b,c) Immunofluorescence staining with the indicated markers and confocal imaging 7 days after TAM injection and 30 min after i.v cadaverine injection. (d) Quantification of brain dye content 7 days after TAM injection and 30min after i.v. injection of dyes with increasing MW (n = 5 mice/group). (e,f) Adult *Unc5B^fl/fl^* and Unc5BiECko brain vessel immunofluorescence using the luminal marker podocalyxin and quantification of vascular density. (g,h) *Unc5B^fl/fl^* was crossed with *PDGFRβCre^ERT2^* and BBB permeability was assessed 7 days after the last TAM injection and 30 min after cadaverine injection i.v. (n > 6 mice/group). (i,j) Western blot and quantification of adult *Unc5B^fl/fl^* and Unc5BiECko brain protein extracts, n = 5 animals/group. (k) Immunofluorescence staining with the indicated antibodies and confocal imaging of adult *Unc5B^fl/fl^* and Unc5BiECko piriform cortex 7 days after TAM injection and 30 min after i.v cadaverine injection. All data are shown as mean+/−SEM. NS: non-significant, PIR: Piriform cortex, RSP: Retrosplenial cortex, HI: Hippocampus, HY: Hypothalamus, TH: Thalamus, ST: Striatum, PTL: Posterior parietal association areas, SSp: Primary somatosensory cortex, CB: Cerebellum, M: Medulla. Mann-Whitney U test was performed for statistical analysis.

Staining of adult brain sections with a commercial antibody recognizing Unc5B showed labeling of endothelium in various brain regions (**Supp. Fig. 2a**) and revealed that Unc5B deletion had no effect on vascular density (**Fig. 1e,f**). Unc5B expression was also detected in a few CD13^+^ pericytes (**Supp. Fig. 2b)**. Because pericytes contribute to BBB integrity^19,20^, we determined whether pericyte-derived Unc5B affected the BBB by crossing the *Unc5B^fl/fl^* mice with *PdgfrβCre^ERT2^* mice^21^ (Unc5BiPCko), to delete *Unc5B* in mural cells. Neither TAM-treated Cre-negative littermate controls, nor Unc5BiPCko mice showed cadaverine leakage across the BBB (**Fig. 1g,h**). Hence, endothelial, but not pericyte, Unc5B controls adult BBB integrity.

### Unc5B regulates Wnt/β-catenin signaling

To determine the cause of the BBB defect, we measured expression levels of Claudin5 and PLVAP as well as markers for pericytes and astrocytes^19,20,22–24^. Compared to TAM-treated Cre-negative littermate controls, Unc5BiECko mice showed significantly reduced expression of Claudin5, along with increased expression of PLVAP (**Fig.1 i,j**). By contrast, western blot and immunostaining showed that expression of other BBB regulators such as Caveolin1 (in ECs), PDGFRβ (in pericytes), GFAP and Aquaporin-4 (in astrocytes) were similar between genotypes (**Supp. Fig. 3 a-d**).

Homozygous global *Unc5B* KO E12.5 embryos^16^ exhibited decreased Claudin5 and increased PLVAP expressions in the brain (**Supp. Fig. 3e-g**), validating *Unc5B* deletion effects on Claudin5 and PLVAP in an independent knockout mouse strain. Immunolabeling of adult brain sections showed decreased Claudin5 and increased PLVAP expression in areas with cadaverine leakage in Unc5BiECko brains (**Fig. 1k**), suggesting that BBB leakage may be due to changes in expression of Claudin5 and PLVAP.

Because Claudin5 and PLVAP are two known targets of Wnt/β-catenin signaling^6–9^, we determined if Unc5B affected Wnt signaling at the BBB. Western-blot analysis on brain lysates revealed a significant decrease of β–catenin and the LEF1 transcriptional effector in Unc5BiECko brains compared to littermate controls (**Fig. 2a-b**), and immunostaining confirmed decreased LEF1 expression in brain ECs of Unc5BiECko mice (**Fig. 2c-d**). Moreover, phosphorylation of LRP6 at S1490, a hallmark of Wnt/β–catenin pathway activation, was dramatically downregulated upon *Unc5B* gene deletion (**Fig. 2a-b**). This phosphorylation provides a docking site for the adapter protein Axin1, resulting in inhibition of the β–catenin destruction complex and thereby promoting β–catenin nuclear translocation and activation^25,26^.

**Figure 2:**
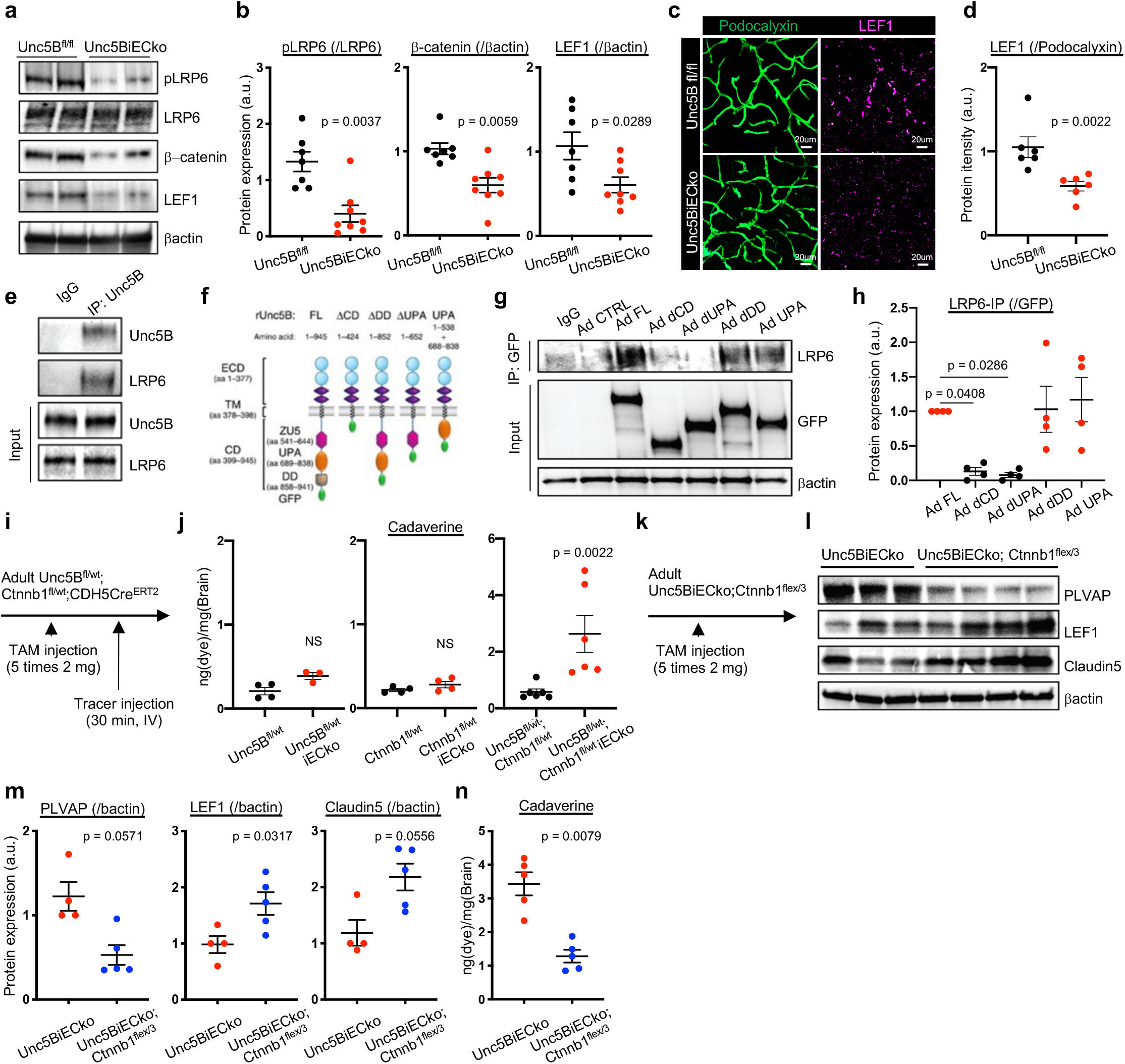
Unc5B regulates the BBB via Wnt/β-catenin signaling. (a,b) Western blot and quantification of Wnt signaling components in adult *Unc5B^fl/fl^* and Unc5BiECko brain protein extracts, n > 7 mice per group. (c,d) Immunofluorescence and quantification of LEF1 staining on adult *Unc5B^fl/fl^* and Unc5BiECko brain vibratome sections. (e) IgG control and Unc5B immunoprecipitation on cultured brain endothelial cells. (f) Unc5B GFP-adenovirus schematic. (g,h) IgG control or GFP immunoprecipitation in ECs infected with Unc5B constructs. (i,j) Quantification of brain cadaverine content in adult mice, 7 days after the last TAM injection and 30 min after i.v cadaverine injection. (k-m) Western blot and quantification of adult Unc5BiECko and Unc5BiECko; Ctnnb1flex/3 brain protein extracts, n > 4 mice per group. (n) Quantification of brain cadaverine content in adult mice, 7 days after the last TAM injection and 30 min after i.v cadaverine injection. (n = 5 mice per group). All data are shown as mean+/− SEM. NS: non-significant. Mann-Whitney U test was performed for statistical analysis between two groups. ANOVA followed by Bonferroni’s multiple comparisons test was performed for statistical analysis between multiple groups.

Immunoprecipitation of Unc5B from primary microvascular mouse brain ECs pulled down LRP6 (**Fig. 2e**), demonstrating a physical interaction between Unc5B and LRP6 receptors. To determine which Unc5B domain mediated this interaction, we infected Unc5B siRNA-treated human ECs with GFP-tagged siRNA resistant rat adenoviral constructs encoding Unc5B full-length (FL) or a cytoplasmic domain deletion (ΔCD) (**Fig. 2f**). LRP6 co-IP was rescued by Unc5B FL but not by ΔCD, identifying the Unc5B cytoplasmic domain as the main LRP6 interacting domain. Additional cytoplasmic domain deletions revealed that the Unc5B death domain (DD) that mediates apoptosis in the absence of the ligand^27,28^ was dispensable for LRP6 interaction, whereas deletion of the UPA domain (named after its conservation in Unc5B, PIDD and Ankyrin^28^) abolished LRP6 co-IP. Finally, a construct encoding only the cytoplasmic UPA domain was sufficient to rescue LRP6 co-IP (**Fig. 2f-h**). Hence, Unc5B interacts with LRP6 via its UPA domain.

To test genetic interaction of Unc5B with Wnt/β–catenin signaling, we crossed Unc5BiECko mice with *Ctnnb1^fl/fl^* mice^29^. Heterozygous Unc5B or β–catenin mice displayed no BBB cadaverine leakage, but double heterozygous Unc5B^fl/wt^; β–catenin^fl/wt^-Cdh5Cre^ERT2^ exhibited BBB cadaverine leakage compared to TAM treated Cre- controls (**Fig. 2i,j**), demonstrating that Unc5B and β–catenin genetically interact to maintain BBB integrity.

Next, we crossed Unc5BiECko with mice overexpressing an activated form of β–catenin (*Ctnnb1^flex/3^* mice) (**Fig. 2k**), which enhances Wnt/β–catenin signaling^30^. The resulting offspring (Unc5BiECko; Ctnnb1^flex/3^) displayed decreased PLVAP protein expression, along with increased LEF1 and Claudin5 protein expression compared to Unc5BiECko mice (**Fig. 2l,m**). Cadaverine injection into Unc5BiECko; Ctnnb1^flex/3^ mice showed that BBB leakage was reduced by β–catenin overexpression in Unc5BiECko mice (**Fig. 2n**).

We considered other signaling pathways that could contribute to BBB leakage in Unc5BiECko mice. Unc5B inhibits Vegfr2-mediated permeability signaling in ECs *in vitro* by reducing phosphorylation of the Y949 residue^13^. Y949 phosphorylation is known to trigger disassembly of adherens junctions by activating VE-Cadherin phosphorylation, which then downregulates Claudin5 ^12,13,31^. Therefore, increased brain Vegfr2-Y949 permeability signaling in the absence of Unc5B could contribute to BBB opening. Western blotting of brain lysates revealed increased Vegfr2-Y949 phosphorylation in Unc5BiECko compared to Cre- littermate controls, while Vegfr2-Y1173 phosphorylation, which is critical for VEGF-induced proliferation, was unaffected (**Supp. Fig. 4a,b**). To test Vegfr2-Y949 function, we crossed Unc5BiECko mice with Vegfr2-Y949F mutant mice, which carry an inactivating substitution of tyrosine to phenylalanine and are resistant to VEGF-induced permeability^32^. Injection of fluorescent cadaverine revealed increased dye leakage into the brain of Unc5BiECko; Y949F mice compared to Cre- littermate controls (**Supp. Fig. 4c,d**), demonstrating that Vegfr2-Y949F failed to rescue BBB integrity in Unc5B mutant mice.

To discriminate between transcellular and paracellular vascular leakage in Unc5BiECko mice, we crossed Unc5BiECko mice with eGFP::Claudin5 mice that express 2-fold higher Claudin5 levels compared to wildtype littermates^33^ and thereby display enhanced paracellular barrier properties of CNS ECs. BBB leakage of Cadaverine into the brains of Unc5BiECko; eGFP::Claudin5 mice was reduced compared to Unc5BiECko mice (**Supp. Fig. 4e,f**), demonstrating that Unc5B may regulate BBB leakage by modulating the levels of Claudin5. However, leakage of 40kDa dextran remained increased in Unc5BiECko; eGFP::Claudin5 mice compared to Unc5BiECko mice (**Supp. Fig. 4g**), suggesting that loss of Unc5B induced both paracellular leak for small MW tracers mediated by loss of Claudin5 as well as transcellular leak for higher MW dyes potentially by inducing PLVAP.

### Netrin1 binding to Unc5B mediates BBB integrity

We then investigated whether Unc5B ligands Netrin1 and Robo4 regulated the Wnt/β–catenin pathway activation in CNS ECs. Since Netrin1 mRNA is produced by several cell types in the adult brain^34^, we generated temporally inducible *Netrin1* global KO mice by crossing *Ntn1^fl/fl^* mice with *RosaCre^ERT2^* mice (hereafter Ntn1iko), to induce ubiquitous gene deletion upon TAM injection. Compared to TAM-treated Cre- littermate controls, i.v. injection of cadaverine in adult Ntn1iko mice revealed increased cadaverine leakage across the BBB (**Fig. 3a**), while *Robo4* KO mice^12^ did not exhibit any BBB deficits (**Fig. 3b**). Further analysis of adult Ntn1iko mouse brain lysates revealed efficient *Ntn1* gene deletion along with decreased pLRP6, Claudin5 and LEF1 protein expression, while PLVAP expression was increased (**Fig. 3c,d**). Moreover, treating serum-starved mouse brain primary ECs with Netrin1 increased LRP6 phosphorylation with a peak at 30min to 8h after stimulation (**Fig. 3e,f**). This effect was abolished by *Unc5B* siRNA treatment (**Fig.3 e,f**). Unc5B immunoprecipitation from mouse brain lysates revealed reduced LRP6 co-IP in the Ntn1iko mice when compared to controls (**Fig. 3g,h**), suggesting that Netrin1 binding to Unc5B regulated LRP6 phosphorylation and Wnt/β–catenin activation in CNS ECs.

**Figure 3:**
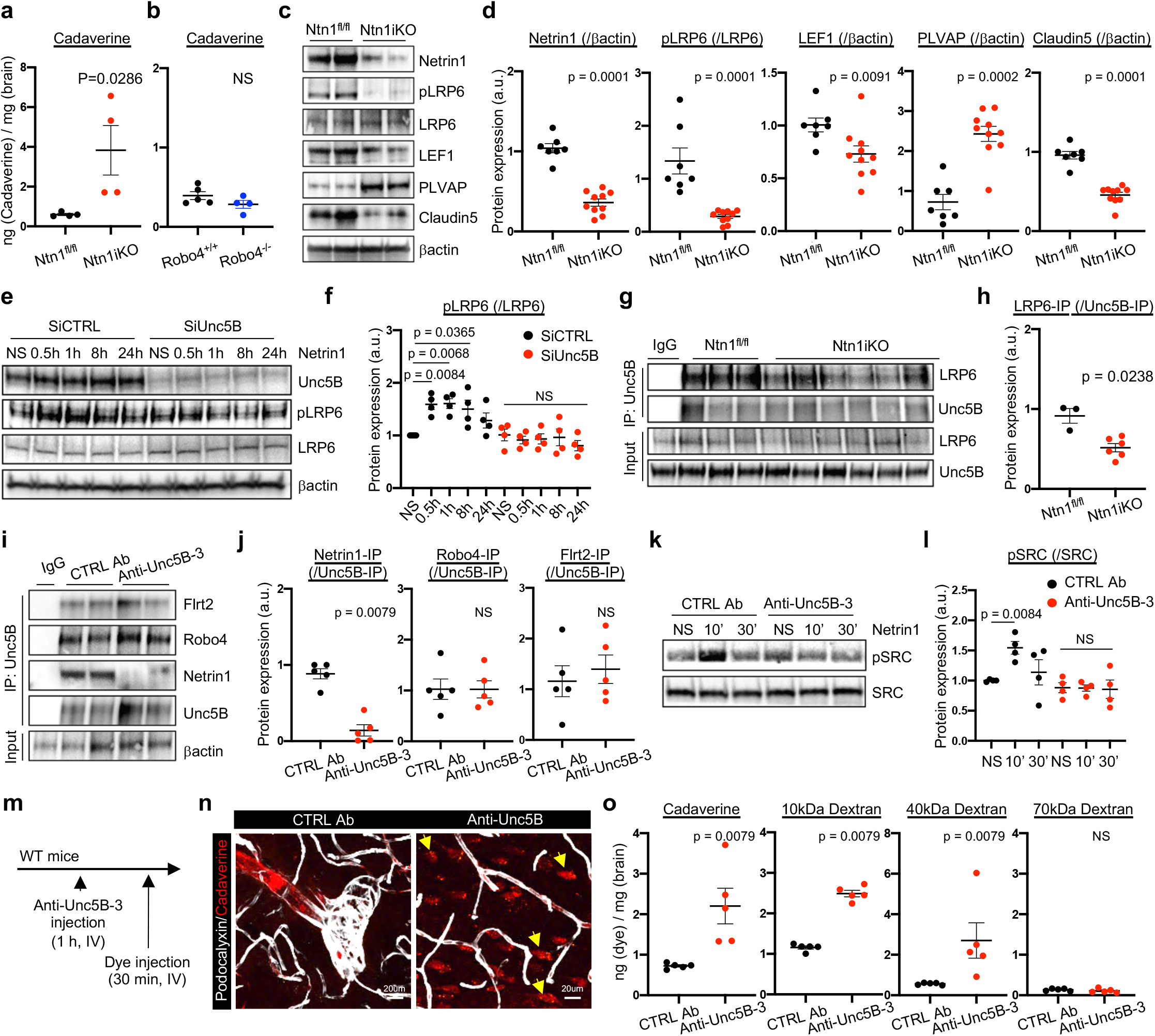
Netrin1 binding to Unc5B regulates Wnt/β-catenin signaling to maintain BBB integrity. (a.b) Quantification of brain cadaverine content in adult mice, 7 days after the last TAM injection (a) and 30 min after i.v cadaverine injection (n = 4 mice/group). (c,d) Western blot and quantification of adult brain protein extracts from *Ntn1^fl/fl^* and Ntn1iKO mice (n>7 mice per group). (e,f) Western blot and quantification of brain ECs treated with SiCTRL or SiUnc5B before Netrin1 treatment for the indicated times. (g,h) Unc5B immunoprecipitation of brain protein extracts and LRP6 western blot and quantification. (i,j) Unc5B immunoprecipitation of brain protein extracts from mice i.v injected with CTRL or anti-Unc5B-3 antibodies (1 h, 10 mg/kg) and western blot with antibodies recognizing the indicated ligands and quantification (n = 5 mice per group). (k,l) Western-blot and protein quantification of ECs treated with CTRL or anti-Unc5B-3 for 1h followed by Netrin1 stimulation (500 ng/ml) for 10 min or 30 min (n = 4). (m,n) BBB permeability was assessed by immunofluorescence on brain vibratome section from mice injected with CTRL or anti-Unc5B-3 antibodies i.v. (1 h, 10 mg/kg). (o) Quantification of brain dye content 30 min after injection of dyes with increasing MW and 1 h after CTRL or anti-Unc5B-3 i.v. injection (10 mg/kg). All data are shown as mean+/−SEM. NS: non-significant. Mann-Whitney U test was performed for statistical analysis between two groups. ANOVA followed by Bonferroni’s multiple comparisons test was performed for statistical analysis between multiple groups.

To specifically interrogate whether blocking Netrin1-Unc5B interactions disrupted the BBB *in vivo*, we used monoclonal antibodies (mAbs) that we had previously generated against the Unc5B IgG-like domains^12^. Anti-Unc5B-1 recognizes human but not mouse Unc5B, while anti-Unc5B-2 recognizes both human and mouse Unc5B and internalizes Unc5B^12^. Anti-Unc5B-2 treatment induced Unc5B internalization in brain ECs *in vitro* (**Supp. Fig. 5a**) and i.v. injection of Anti-Unc5B-2 for 1 hour at 10mg/kg in mice reduced brain Unc5B expression compared to anti-Unc5B-1 CTRL Ab-treated animals (**Supp. Fig. 5b,c**), therefore preventing binding of all Unc5B ligands *in vivo*.

To generate a mAb that specifically blocked Netrin1 binding without Unc5B internalization, we screened a human phage-derived library against the entire rat Unc5B extracellular domain and identified anti-Unc5B-3, a mAb that bound both human and mouse Unc5B with high affinity (**Supp. Fig. 5d-f**) but did not induce Unc5B internalization nor its degradation *in vivo* (**Supp. Fig. 5g,i**). I.v. injection of anti-Unc5B-3 for 15min at 10mg/kg followed by cardiac perfusion and immunolabelling using an anti-human IgG antibody revealed anti-Unc5B-3 binding to the brain vasculature of *Unc5B^fl/fl^*, but no binding in the Unc5BiECko mice (**Supp. Fig. 5j,k**), demonstrating specific binding of anti-Unc5B-3 to endothelial Unc5B. I.v. injection of anti-Unc5B-3 (10mg/kg for 1h) followed by Unc5B immunoprecipitation revealed that anti-Unc5B-3 blocked Netrin1 binding to Unc5B *in vivo* compared to CTRL Ab treated mice, while Robo4 and Flrt2 could still interact with Unc5B (**Fig. 3i,j**). Anti-Unc5B-3 also blocked Netrin1-induced Src phosphorylation in brain ECs *in vitro* (**Fig. 3k,l**).

### Transient ‘on demand’ BBB opening via anti-Unc5B antibodies

To test if antibody mediated Unc5B blockade could be used to open the BBB “on-demand”, we injected i.v. CTRL or anti-Unc5B antibodies for 1hr in adult WT C57BL/6J mice, followed by i.v. injection of fluorescent tracers of various molecular weights 30min before sacrifice and analysis (**Fig. 3m**). In mice treated with CTRL Ab, there were no signs of BBB disruption and injected cadaverine remained confined inside brain vessels (**Fig. 3n,o**). In contrast, mice treated with anti-Unc5B-3 or anti-Unc5B-2 showed a significant leakage of injected Cadaverine, 10kDa and 40kDa Dextran, but not 70kda Dextran or endogenous immunoglobulin or fibrinogen into the brain parenchyma (**Fig. 3n,o** and **Supp. Fig. 6a-c**), demonstrating that blocking Netrin1 binding to Unc5B is sufficient to open the BBB. The vascular barrier disrupting effects of anti-Unc5B-2 and -3 were specific to the brain, as tracer leakage in other organs was similar between controls and anti-Unc5B-2 or -3 treated mice (**Supp. Fig. 6d,e**).

Anti-Unc5B-3 treatment also enhanced delivery of single chain nanobodies across the BBB when compared to CTRL Ab (**Fig.4a**), while nanobody extravasation in other organs such as lung, heart, kidney or skin remained similar (**Supp. Fig. 6f**). Moreover, injection of anti-Unc5B-3 mAbs enhanced brain delivery of BDNF and induced phosphorylation and activation of its neuronal receptor Trk-B, while plasma BDNF levels remained similar to CTRL Ab injected mice (**Fig.4b-d**), indicating that bioactive molecules up to 40kDa can be delivered into the brain by this approach.

**Figure 4:**
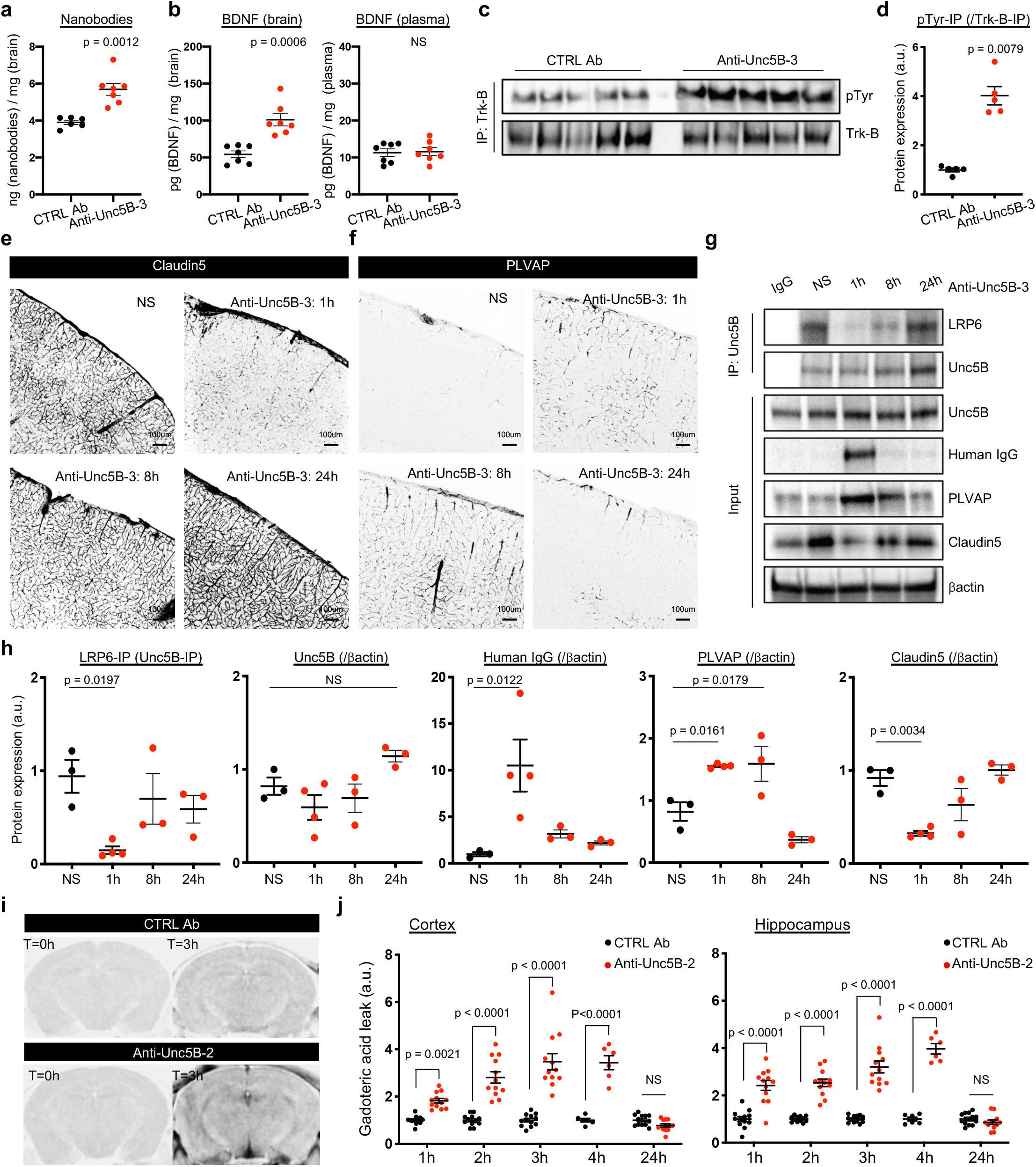
Reversible BBB opening by Unc5B-blocking antibodies. (a) Quantification of brain nanobody content 1h after i.v CTRL or anti-Unc5B-3 injection (10 mg/kg) and 30min after i.v nanobody injection. (b) Quantification of brain and plasma BDNF concentration 1 h after i.v CTRL or anti-Unc5B-3 injection (10mg/kg) and 30m in after i.v BDNF injection. (c,d) Trk-B immunoprecipitation and anti-phospho-tyrosine western blot on brain protein extracts from mice injected with CTRL or anti-Unc5B-3 antibodies i.v. (1h, 10mg/kg) followed by BDNF injection for 30 min (n = 5 mice per group). (e,f) Immunofluorescence staining of Claudin5 and PLVAP on adult brain vibratome section from mice i.v. injected with anti-Unc5B-3 (10 mg/kg) for 1, h or 24 h. (g,h) Unc5B immunoprecipitation and quantification of brain protein extracts from mice injected with anti-Unc5B-3 i.v. (10 mg/kg) for 1, 8 or 24 h, n = 3/4 animals/group. (i,j) MRI analysis and quantification of gadolinium leakage after CTRL or anti-Unc5B-2 injection. Gadolinium was injected 1, 2, 3, 4 and 24 h after the antibodies. All data are shown as mean+/−SEM. NS: non-significant. Mann-Whitney U test was performed for statistical analysis between two groups. ANOVA followed by Bonferroni’s multiple comparisons test was performed for statistical analysis between multiple groups.

To determine the specific BBB vascular beds regulated by Unc5B, we injected TAM in adult *BMXCre^ERT2^-mTmG* mice that specifically express GFP in arteries upon TAM injection, followed by anti-Unc5B-3 i.v. injection for 15 min and cardiac perfusion. Human IgG staining of anti-Unc5B-3 revealed binding to GFP+ brain arteries at cell-cell junctions but also to GFP-capillaries (**Supp. Fig.7a**), suggesting that Unc5B regulates BBB integrity mainly in arteries and capillaries.

To assess anti-Unc5B bioavailability and vascular clearance, we injected anti-Unc5B-3 antibodies (10mg/kg) i.v. for 1h, 8h or 24h followed by immune-labeling with anti-human IgG antibodies (**Supp. Fig.7b**). Anti-Unc5B-3 was detectable in the brain vasculature 1h after injection, declined to low levels after 8h and was undetectable 24h after injection (**Supp. Fig.7b**), demonstrating rapid clearance from the brain vasculature.

Interestingly, the expression of Wnt/β–catenin downstream targets varied in a similar time-dependent fashion. Claudin5 immunostaining was downregulated 1h after anti-Unc5B-3 injection and returned to basal levels after 8h, whereas PLVAP immunostaining was upregulated at 1 and 8h after anti-Unc5B-3 injection and returned to low baseline levels after 24h (**Fig. 4e,f**). Western blot on brain protein lysates from mice treated with CTRL Ab or anti-Unc5B-3 for 1h, 8h or 24h confirmed these changes (**Fig. 4g,h**). Unc5B immunoprecipitation showed that anti-Unc5B-3 treatment transiently disrupted the Unc5B/LRP6 interaction 1h after anti-Unc5B-3 injection, thereby disrupting Wnt/β–catenin signal transduction (**Fig. 4g,h**).

Finally, we provided a real time evidence of the transient opening of the BBB mediated by anti-Unc5B treatment using both MRI and two-photon live imaging through cranial windows. Mice received i.v. injection of CTRL or anti-Unc5B-2 mAb and were imaged by contrast MRI after i.v. injection of gadoteric acid (MW 558 Da) at 1, 2, 3,4 and 24 hours following antibody treatment. Quantification showed a significant gadoteric acid leakage in the cortex and hippocampus between 1-4 hours after anti-Unc5B-2 delivery, which returned to baseline levels after 24 hours (**Fig. 4i,j**), indicating that the neurovascular barrier had resealed. Two-photon live imaging showed that a 2000kDa FITC-dextran did not leak at any time point after anti-Unc5B-2 treatment but outlined the brain vasculature (**Supp. Fig. 8a**). By contrast, Hoechst (MW 560Da) started to extravasate from superficial cortical vessels within 5min after injection and continued to leak over 30 minutes in mice that were treated with the anti-Unc5B-2 antibody one hour prior to tracer injection (**Supp. Fig. 8a**). When 10kDa dextran was injected intravenously in mice that were treated with the anti-Unc5B-2 antibody 24 hours earlier, we did not observe any tracer leakage (**Supp. Fig. 8b**), indicating that the BBB had resealed. However, re-administration of a second dose of anti-Unc5B-2 re-opened the BBB within an hour leading to 10kDa dextran extravasation over the next 30 min (**Supp. Fig. 8b**).

## Discussion

In summary, our data reveal Netrin1 signaling to Unc5B as a novel BBB regulatory pathway with potential therapeutic relevance in CNS disease (**Supp. Fig. 9**). We showed that mice deficient in either endothelial Unc5B receptor or Netrin1 ligand exhibited BBB leakage accompanied by reduced expression of Wnt/β-catenin components, β-catenin and LEF1. Moreover, i.v. delivery of Unc5B mAbs that specifically block Netrin1 binding to Unc5B (anti-Unc5B-3), or that block binding of all Unc5B ligands via receptor internalization (anti-Unc5B-2), led to transient Wnt/β-catenin signaling reduction and BBB breakdown, supporting that Netrin1 binding to Unc5B is sufficient to maintain Wnt signaling activation in CNS ECs and BBB integrity. Further experiments are required to determine the source of Netrin1 mediating this effect. Single cell RNA sequencing studies indicate that Netrin1 is highly expressed in adult brain pericytes, indicating that pericytes are a likely source of Netrin1 production and Unc5B activation at the BBB^34^. Interestingly, Netrin1 was shown to be implicated in BBB integrity, upregulated in brain ECs in response to brain injury, and to increase Claudin5 expression^35,36^, therefore multiple cellular sources and environmental modulations of Netrin1 expression could contribute to BBB integrity. Since we could target Unc5B via i.v. blocking antibody injection, these data raise the possibility that i.v. injection of Netrin1 or other Unc5B agonists could repair CNS endothelial barrier breakdown in conditions such as ischemic stroke or multiple sclerosis where the BBB is dysfunctional.

The BBB leakage in *Unc5B* mutants was region-specific and affected caudal and ventral regions more than anterior and dorsal ones, which is roughly similar to the region-specific BBB leakage observed in young adult mice carrying inducible allelic deletions of β-catenin^9,37^. The specific Wnt ligands and receptors that maintain the BBB differ in a region-specific manner in the CNS, with cerebellum BBB utilizing Norrin, LRP5/6 and TSPAN12 signaling module, while cortex relies on Wnt7a/b, GPR124, and RECK. Ligands and receptors in different brain regions are partially redundant, in that inactivation of several components exacerbates BBB leakage^5,9,37^. Remarkably, blockade of Unc5B function affected both the cortex, cerebellum and other brain regions, suggesting that Unc5B may be an upstream regulator of several Wnt/Norrin signaling complexes at the BBB. Mechanistically, we identify the Unc5B intracellular UPA domain as a regulator of LRP6 interaction, suggesting that the UPA domain may induce LRP6 phosphorylation through recruitment of kinases or other mechanisms that remain to be determined. Recent studies in naïve pluripotent embryonic stem cells showed that Netrin1 binding to Unc5B induced FAK-mediated phosphorylation of GSK3α/β, a kinase implicated in LRP6 activation ^38^, suggesting one possible mechanism. Finally, because Unc5B is expressed in arterial and capillary endothelium, but not in veins, it is likely to confer BBB integrity in a vessel-segment specific manner, underscoring heterogeneity of BBB regulation in different vascular segments^39^.

Previous studies had speculated that transient Wnt signaling inhibition could be used to open the BBB “on-demand” for drug delivery into the diseased CNS^9,40^, but the means to inhibit Wnt signaling in a CNS specific manner were not available. We demonstrated that antibody mediated Unc5B blockade caused a transient loss of Wnt/β–catenin signaling and BBB breakdown for 1h to 8h followed by neurovascular barrier resealing, and allowed delivery of tracers up to 40kDa into the adult CNS. The size selectivity of BBB opening is compatible with delivery of chemotherapeutics and of bioactive molecules such as nanobodies and growth factors. Anti-Unc5B mAbs could therefore synergize with existing therapies such as focused ultrasound/microbubble approaches^41–44^ and offer a new therapeutic perspective for treatment of various human neurological disorders.

## Methods

### Mouse models

All protocols and experimental procedures were approved by the Institutional Animal Care and Use Committee (IACUC). Generation of the targeted Unc5b allele was performed by homologous recombination in R1 ES cells. Correctly targeted cells were identified by Southern blot hybridization and injected into B6J blastocysts to generate Unc5bneo/+ mice. To remove the neo cassette, Unc5bneo/+ mice were mated to B6.129S4-Gt(ROSA)26Sortm1(FLP1)Dym/RainJ mice (The Jackson Laboratory, stock #009086). Mice were backcrossed to B6J mice for ten generations. Unc5B^fl/fl^ (B6-Unc5b<tm1(flox)Slac/Slac) mice were then bred with Cdh5Cre^ERT2^ mice^17^ or PDGFRβCre^ERT2 21^. eGFP::Claudin5 transgenic mice, Y949F mice, β–catenin GOF Ctnnb1^flex/3^ mice, β–catenin^fl/fl^ mice, Robo4^−/−^ mice and Netrin1^fl/fl^ mice were described previously^12,29,30,32,33,45^. Gene deletion was induced by injection of tamoxifen (Sigma T5648) diluted in corn oil (Sigma C8267). Postnatal gene deletion was induced by 3 injections of 100ug of tamoxifen at P0, P1 and P2; whereas adult gene deletion was induced by 5 injections of 2mg of tamoxifen from P60 to P64.

### Cell culture

Bend3 cells were purchased from ATCC (ATCC® CRL-2299™) and C57BL/6 Mouse Primary Brain Microvascular Endothelial Cells were purchased from Cell Biologics (C57-6023). Cells were cultured in Dulbecco’s Modified Eagle’s Medium (DMEM) high glucose (Thermo Fisher Scientific, 11965092) supplemented with 10% fetal bovine serum (FBS) and 1% Penicillin Streptomycin at 37 °C and 5% CO2 and split when confluent using Trypsin-EDTA (0.05%) (Life Technologies, 25300054). When indicated, cells were stimulated using Recombinant Mouse Netrin-1 Protein (R&D, 1109-N1-025) at 500ng/ml.

### Western-Blot

Brains were dissected and frozen in liquid nitrogen. They were lysed in RIPA buffer (Research products, R26200-250.0) supplemented with protease and phosphatase inhibitor cocktails (Roche, 11836170001 and 4906845001) using a Tissue-Lyser (5 times 5min at 30 shakes/second). For western blot on cell culture, cells were washed with PBS and lysed in RIPA buffer with protease and phosphatase inhibitors cocktails. All protein lysates were then centrifuged 15min at 13200RPM at 4°C and supernatants were isolated. Protein concentration were quantified by BCA assay (Thermo Scientific, 23225) according to the manufacturer’s instructions. 30ug of protein were diluted in Laemmli buffer (Bio-Rad, 1610747) boiled at 95°C for 5min and loaded in 4-15% acrylamide gels (Bio-Rad, 5678084). After electrophoresis, proteins were transferred on a polyvinylidene difluoride (PVDF) membrane and incubated in TBS 0.1% Tween supplemented with 5% BSA for 1hour to block non-specific binding. The following antibodies were incubated overnight at 4°C: Unc5B (Cell Signaling, 13851S), Robo4 (Invitrogen, 20221-1-AP), Flrt2 (Novus bio, NBP2-43653), Netrin1 (R&D, AF1109), Claudin5 (Invitrogen, 35-2500), PLVAP (BD biosciences, 550563), PDGFRβ (Cell Signaling, 3169S), GFAP (DAKO, Z0334), VEGFR2 Y949 (Cell Signaling, 4991S), VEGFR2 Y1173 (Cell Signaling, 2478S), pLRP6 (Cell Signaling, 2568S), LRP6 (Cell Signaling, 3395S), β–catenin (Cell Signaling, 8480S), LEF1 (Cell Signaling, 2230S) and βactin (Sigma, A1978). Then, membranes were washed 4 × 10min in TBS 0.1% Tween and incubated with one of the following peroxidase-conjugated secondary antibodies diluted in TBS 0.1% Tween supplemented with 5%BSA for 2h at room temperature: horse anti-mouse IgG(H+L) (Vector laboratories, PI-2000), goat anti-rabbit IgG(H+L) (Vector laboratories, PI-1000), goat anti-rat IgG(H+L) (Vector laboratories, PI-9400), horse anti-goat IgG(H+L) (Vector laboratories, PI-9500). After 4 × 10min wash, western blot bands were acquired using ECL western blotting system (Thermo Scientific, 32106) or west femto maximum sensitivity substrate (Thermo Scientific, 34095) on a Biorad Gel Doc EQ System with Universal Hood II imaging system equipped with Image Lab software.

### Immunoprecipitation

Pierce™ protein A/G magnetic beads (Thermo fischer, 88802) were washed 5 times 10min with RIPA buffer. 300ug of protein lysate were diluted in 1ml of RIPA buffer containing protease and phosphatase inhibitors and were incubated with 30ul of A/G magnetic beads for 1hour at 4°C under gentle rotation. Protein lysates were harvested using a magnetic separator (Invitrogen) and were incubated overnight at 4°C under gentle rotation with 10ug of Unc5B antibody (R&D, AF1006) or control IgG. The next day, 40ul of protein A/G magnetic beads were added to each protein lysate for 2hour at 4°C under gentle rotation. Beads were then isolated using a magnetic separator and washed 5 x with RIPA buffer. After the last wash, supernatants were removed and beads were resuspended in 40ul of Laemmli buffer (Bio-Rad, 1610747), boiled at 95°C for 5min and loaded onto 4-15% gradient acrylamide gels. Western blotting was performed as described above.

### Immunostaining

Brains were collected and placed in 3.7% formaldehyde overnight at 4°C. Brains were then washed 3 times 10min with TNT buffer (for 100ml: 10ml Tris 1M pH7,4, 3ml NaCl 5M, 500ul Triton X-100) and embedded in 2% agarose. 150um sections were prepared using a Leica VT 1000S vibratome and placed in TNTB buffer (TNT buffer supplemented with 5% donkey serum) for 24h at 4°C. Primary antibodies were diluted in TNTB and placed for 48h at 4°C under gentle agitation. Then, sections were washed 5 times 30min with TNT buffer and incubated for 24h at 4°C with secondary antibodies diluted in TNTB buffer. After 5 × 30min wash with TNT, sections were mounted using DAKO mounting medium (Agilent, S302380-2).

For brain endothelial cell immunostaining, cells were seeded on 18mm glass coverslips (Fischer Scientific, 12542A). When confluent, cells were washed with PBS and fixed with 3.7% formaldehyde for 10min. Cells were washed 3 times 5min with PBS and were incubated with 0.2% TritonX100 diluted in PBS for an additional 10min, washed 3 times and incubated with blocking solution (2%BSA, 3%Donkey serum diluted in PBS) for 1hour at room temperature. Primary antibodies were then diluted in blocking solution and incubated on coverslips overnight at 4°C. After 3 × 5min washes, secondary antibodies diluted in blocking buffer were incubated on coverslips for 2h at room temperature. Coverslips were then washed 3 times 5min with PBS and mounted using DAKO mounting medium.

The following antibodies were used: Podocalyxin (RD, AF1556), Unc5B (Cell signaling, 13851S), Claudin5-GFP (Invitrogen, 352588), GFAP (Millipore, MAB360), Aquaporin4 (Millipore, AB3068), PDGFRβ (Cell Signaling, 3169S), LEF1 (Cell Signaling, 2230S), Endomucin (Hycult biotech, HM1108), fibrinogen (DAKO, A0080), DAPI (Thermo Fischer, 62248). All corresponding secondary antibodies were purchased from Invitrogen as Alexa Fluor (488, 568, 647) donkey anti-primary antibodies (H+L).

### Small interfering RNA knockdown experiments

For Unc5B Inhibition, cells were transiently transfected with siRNA (Dharmacon). ON-TARGETplus Mouse Unc5b siRNA (SMARTpool, L-050737-01-0005) were used for Unc5B gene deletion. Transfection was performed using lipofectamine RNAi max (Invitrogen, 13778-075) according to the manufacturer’s instruction with siRNA at a final concentration of 25pmol in OptiMem for 8h. After transfection, cells were washed with PBS and fresh complete media was added for 48h.

### Mouse lung endothelial cell isolation

Mouse lung were collected and minced into small pieces. Lungs were incubated in digestion buffer (5ml of DMEM supplemented with 5mg of collagenase I (Worthington LS004196), 10ul of 1M Ca2+ and 10ul of 1M Mg2+) for 1hour at 37°C with shaking every 10min. Once fully lysed, lung lysates were filtered through a 40um cell strainer (Falcon, 352340) into a solution of 3ml FBS. Samples were centrifuged for 10min at 1500RPM and pellets were resuspended in PBS 0.1%BSA. In the meantime, rat anti-mouse CD31 (BD Pharmigen, 553370) was incubated with sheep anti-rat IgG magnetic dynabeads (Invitrogen, 11035) in a solution of sterile PBS 0.1%BSA (120ul of beads, 24ul of antibodies in 12ml PBS 0.1%BSA). Solutions were place under gentle rotation at room temperature for 2hours to allow proper coupling of antibodies and beads. Coupled beads were next isolated using a magnetic separator and incubated in the resuspended lung lysate for 30min. After 5 washes with PBS 0.1%BSA, beads were separated using magnetic separator and seeded in 60mm dishes containing mouse lung endothelial cell media (DMEM high glucose, 20%FBS, 1% Penicillin Streptomycin, 2% mitogen (Alta Aesar BT203). Purified endothelial cells were cultured at 37 °C and 5% CO2 until confluence was reached, and then harvested.

### Quantitative real-time PCR analysis

mRNA were isolated using Trizol reagent (Life Technologies, 15596018) according to the manufacturer’s instructions and quantified RNA concentrations using nanodrop 2000 (Thermo Scientific). 300ng of RNA were reverse transcribed into cDNA using iScript cDNA synthesis kit (Bio-rad, 170-8891). Real-time qPCR was then performed in duplicates using CFX-96 real time PCR device (Bio-rad). Mouse GAPDH (QT01658692) was used as housekeeping gene for all reactions.

### Unc5B function blocking antibody generation

We performed a Phage-Fab (antigen-binding fragment) selection using a naïve Fab library (libF^46^) on an immobilized recombinant rat Unc5B-ECD Fc fusion protein (R&D systems). Phage particles fused with Fabs were incubated with an unrelated protein (e.g. streptavidin) immobilized on a solid surface and allowed to bind in a step termed counterselection to remove unwanted phage-Fabs prior to incubation against target. After washing away unbound phage-Fab, phage were eluted from the target and amplified overnight for subsequent rounds. After 5 rounds of this process individual clones from rounds 3-5 were grown in 96-well format and tested by ELISA for their ability to bind antigen specifically. We selected several unique and different positive Fab over 5 rounds of selection, which were subcloned before antibody production (Proteogenix, Schiltigheim, France).

### Surface Plasmon Resonance

Binding of anti-Unc5B antibodies to Human or Rat Unc5B was performed using a Biacore™ 8K (Proteogenix, Schiltigheim, France). Human or Rat Unc5B-ECD-Fc (R&D Systems) were immobilized on a CM5 sensor chip. Each antibody was diluted to gradient concentrations (50nM, 25nM, 12.5nM, 6.25nM, 3.125nM) and flow through CM5 chip. The kinetic parameter was calculated using Bia-evaluation analysis software.

### Intravenous injection of antibodies, fluorescent tracer and nanobodies

CTRL Ab, anti-Unc5B-2 and -3 were injected intravenously into the lateral tail-vein of 8 weeks old adult mice at a concentration of 10mg of antibodies/kg of mice and left to circulate from 1h to 24h depending on the experiment. All fluorescent tracers were injected intravenously into the lateral tail vein of 8 weeks old adult mice and left to circulate for 30min. Lysine-fixable Cadaverine conjugated to Alexa Fluor-555 (Invitrogen) was injected at a concentration of 100ug Cadaverine/20g of mice. Lysine-fixable 10, 40 or 70 kDa dextran conjugated to tetramethylrhodamine (Invitrogen) were injected at a concentration of 250ug dextran/20g of mice. Nanobodies (Alexa Fluor-488 coupled anti-mouse nanobodies, Abnova) were injected at a concentration of 60ug nanobodies/20g of mice and left to circulate for 30min. For postnatal experiment, cadaverine was injected intraperitoneally into the P5 neonates at a concentration of 250ug cadaverine/20g of pups and left to circulates 2h

### Fluorescent tracer and nanobodies extravasation quantification

To assess tracer leak, animals were perfused in the left ventricle with PBS. Brains (and other organs) were then collected, and their weight measured. Next, brains (and other organs) were incubated in formamide (Sigma-Aldrich, F7503) at 56°C for 48hours. Dye fluorescence was then measured using a spectrophotometer at the adequate emission and excitation wavelength. Dye extravasation from Unc5B^fl/fl^, Unc5BiECko, WT treated with CTRL Ab and anti-Unc5B-2 were performed at the Yale Cardiovascular Research Center (New Haven, CT, USA) on a BioTek synergy2 spectrophotometer. Dye extravasation from WT treated with CTRL IgG2b and anti-Unc5B-3 were performed at the Paris Cardiovascular Research Center (Paris, France) on a Flexstation3 spectrophotometer. All results were normalized to the corresponding brain weight and reported to a standard made of known concentrations of dye diluted in formamide. Results are shown as ng of dye per mg of corresponding organ or tissue.

### BDNF extravasation quantification

To assess BDNF leak, 1 hour after antibody injection, 50 μg of Human BDNF diluted in saline was injected intravenously into the lateral tail vein in adult mice and left to circulate for 30min. After sampling whole blood in EDTA-coated tubes, animals were perfused in the left ventricle with saline. Brains were then collected, and their weight measured. Blood was centrifuged at 1,000g for 15 minutes at 4°C, and plasma was then stored at −20°C until use. Dissected brains were frozen in liquid nitrogen. They were lysed in RIPA buffer (Thermo Fisher) supplemented with protease and phosphatase inhibitor cocktails (Roche, 11836170001 and 4906845001) with increasing needle gauges and sonicated (3 times of 3 minutes each). All protein lysates were then centrifuged 15min at 14,000g at 4°C and supernatants were isolated.

BDNF concentration in the plasma and brain lysates were quantified using a DuoSet BDNF ELISA Kit (R&D Systems, DY248) according to manufacturer instructions. Results were normalized to the corresponding brain weight. Results are shown as ng of nanobodies per mg of brain tissue.

### MRI

Magnetic resonance imaging (MRI) was performed in mice under isoflurane anesthesia (2% in air) in a 4.7 T magnetic resonance scanner (Bruker BioSpec 47/40USR). Brain images were obtained using a Spin-Echo (SE) T1 weighted sequence (TE/TR: 15/250 ms; matrix: 128×128; slice thickness: 1 mm; with no gap; 12 averages) in the axial and coronal planes after intravenous injections of 100 μL gadoteric acid (0.1 mmol/mL). Imaging was repeated every hour during the first 4 hours and at 24 hours after antibody injection.

### Two photon Live imaging

Craniotomy was performed by drilling a 5-mm circle between lambdoid, sagittal, and coronal sutures of the skull on ketamine/xylazine anesthetized ROSAmT/mG mice. After skull removal, the cortex was sealed with a glass coverslip cemented on top of the mouse skull. Live imaging was done 2 weeks later. For multiphoton excitation of endogenous fluorophores and injected dyes, we used a Leica SP8 DIVE in vivo imaging system equipped with 4tune spectral external hybrid detectors and an InSightX3 laser (SpectraPhysics). The microscope was equipped with in house designed mouse holding platform for intravital imaging (stereotactic frame, Narishige; gas anesthesia and body temperature monitoring/control, Minerve). Mice were injected intravenously with 10mg/kg of UNC5B blocking or control antibodies and 1 hour later with dextran and/or Hoechst, followed by imaging every five minutes over 30 to 90 minutes.

### Confocal microscopy

Confocal images were acquired on a laser scanning fluorescence microscope (Leica SP8 and Zeiss LSM800) using the appropriate software (LASX or ZEN system). 10X, 20X and 63X oil immersion objectives were used for acquisition using selective laser excitation (405, 488, 547, or 647 nm).

### Statistical analysis

All *in vivo* experiments were done on littermates with similar body weight per condition and reproduced on at least 3 different litters. Statistical analysis was performed using GraphPad Prism 8 software. Mann-Whitney U test was performed for statistical analysis on two groups. ANOVA followed by Bonferroni’s multiple comparisons test was performed for statistical analysis between 3 or more groups.

## Acknowledgements

This project has received funding from the NIH (1R01HL149343-01, 1R01DK120373-01A1 to AE, 5R01NS35900 to SLA), the Leducq Foundation (TNE ATTRACT, AE, LCW), INSERM and the European Research Council (ERC) (grant agreement No. 834161 to AE). KB was supported by a fellowship from the AHA (18POST34070109). We thank Max Thomas for technical assistance. SLA is an investigator of the Howard Hughes Medical Institute.

## Data availability

All data generated are included in this article (main or supplementary information files). Additional information can be obtained from the corresponding author upon reasonable request.

---

**Supplemental Figure 1:**
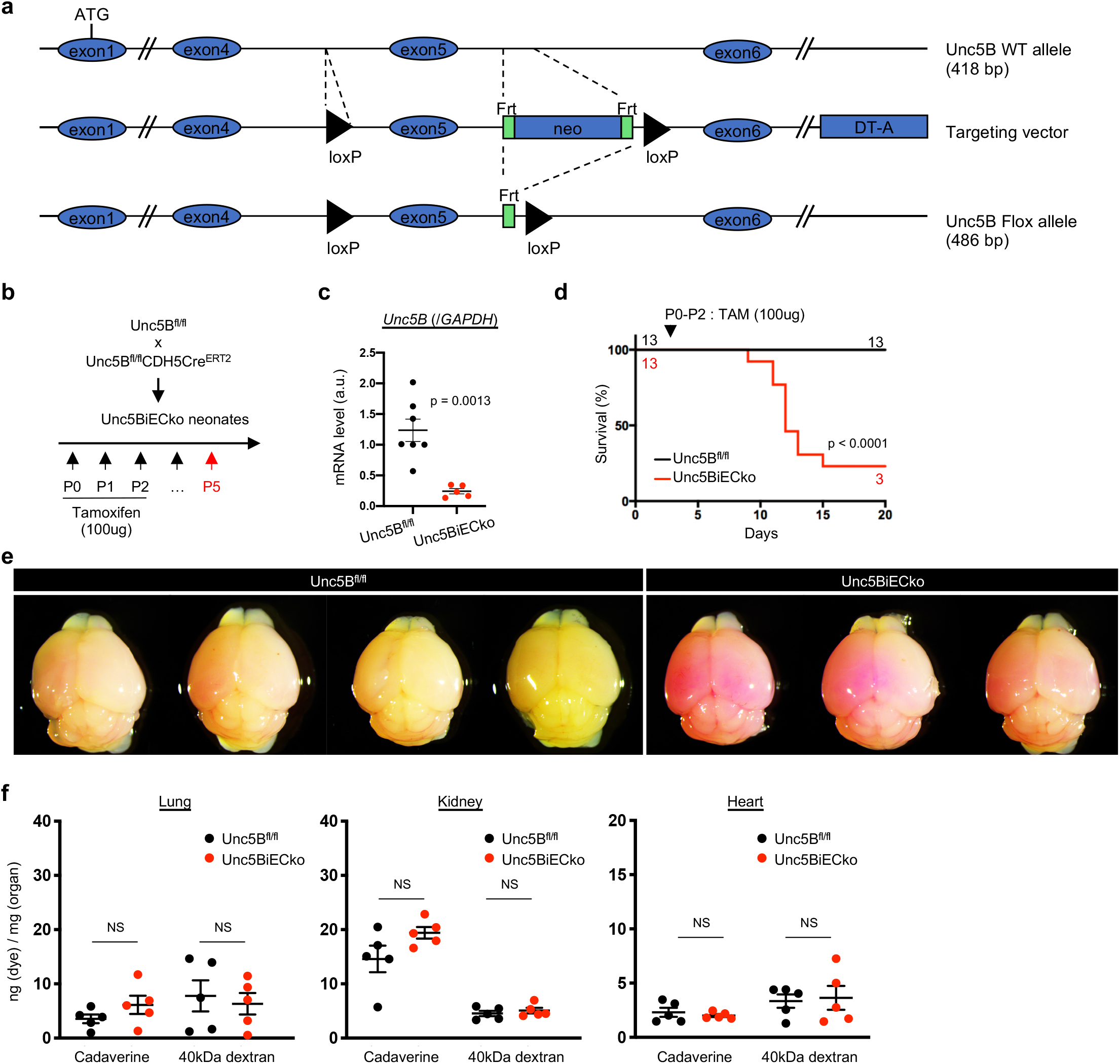
(a) Diagram illustrating generation of the *Unc5B* Flox allele. (b) *Unc5B* gene deletion strategy using tamoxifen (TAM) injection in postnatal mice. (c) qPCR analysis of *Unc5B* mRNA on isolated P5 mouse lung endothelial cells, n>5 mice/group. (d) Survival curve after neonatal *Unc5B* gene deletion (n = 13 mice per group). (e) Cadaverine leakage in P5 *Unc5B^fl/fl^* and Unc5BiECko brains 2 h after intraperitoneal cadaverine injection. (f) Quantification of organ dye content in adult mice, 7 days after the last TAM injection and 30min after i.v injection of dyes with increasing MW (n = 5 mice per group). All data are shown as mean+/−SEM. NS: non-significant. Mann-Whitney U test was performed for statistical analysis between two groups.

**Supplemental Figure 2:**
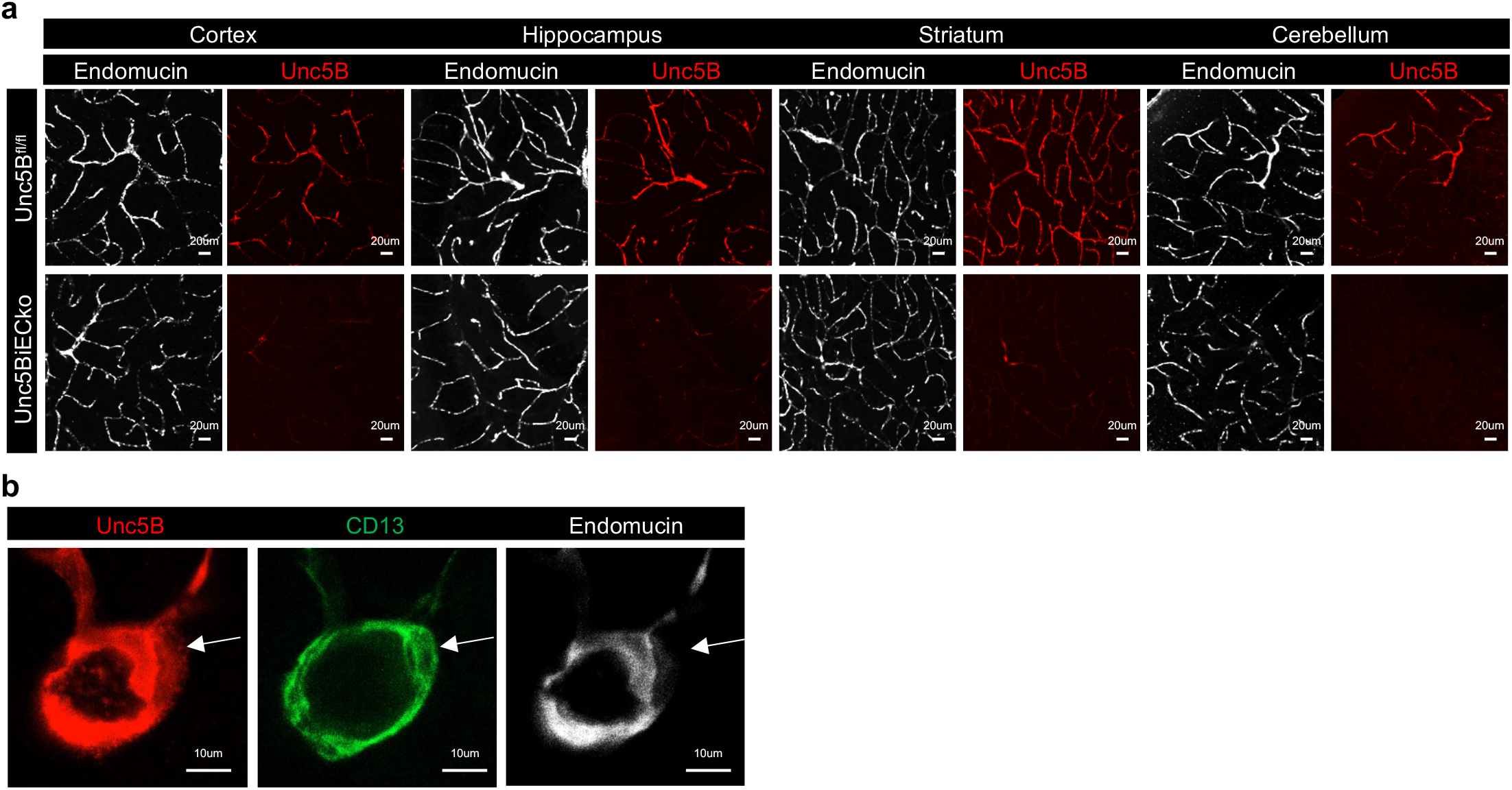
(a,b) Immunofluorescence staining with the indicated antibodies and confocal imaging of vibratome sections from several brain regions.

**Supplemental Figure 3:**
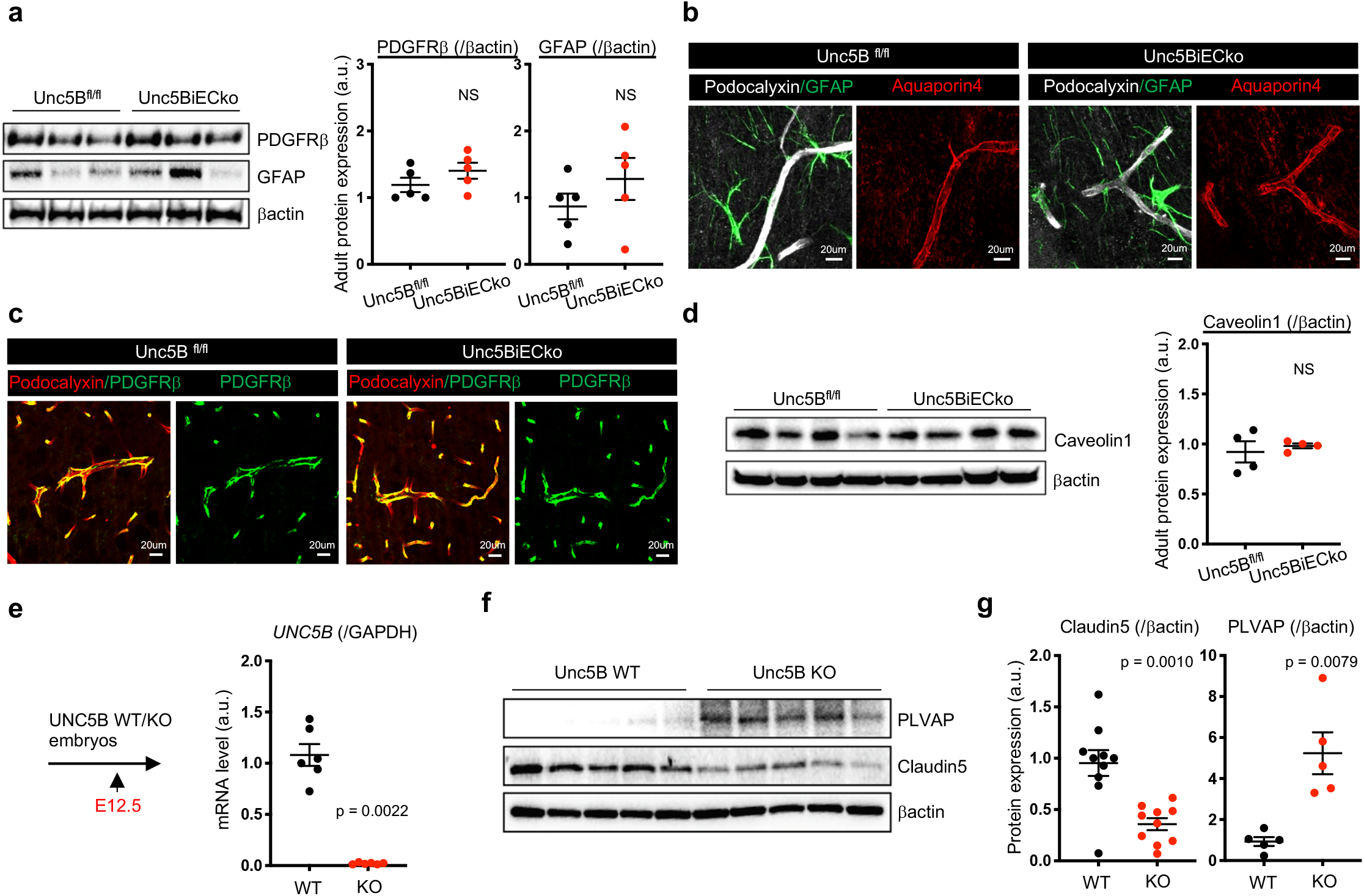
(a) Western blot and quantification of adult *Unc5B^fl/fl^* and Unc5BiECko brain protein extracts (n=5 mice per group). (b,c) Immunofluorescence staining with the indicated antibodies and confocal imaging 7 days after TAM injection. (d) Western blot and quantification of adult *Unc5B^fl/fl^* and Unc5BiECko brain protein extracts, n = 5 animals/group. (e) qPCR analysis on E12.5 brain mRNA extracts from *Unc5B* global KO mice, n = 6 embryos/group. (f,g) Western blot and quantification of E12.5 *Unc5B* WT and KO brain protein extracts (n>5 animals/group). All data are shown as mean+/−SEM. NS: non-significant, Mann-Whitney U test was performed for statistical analysis.

**Supplemental Figure 4:**
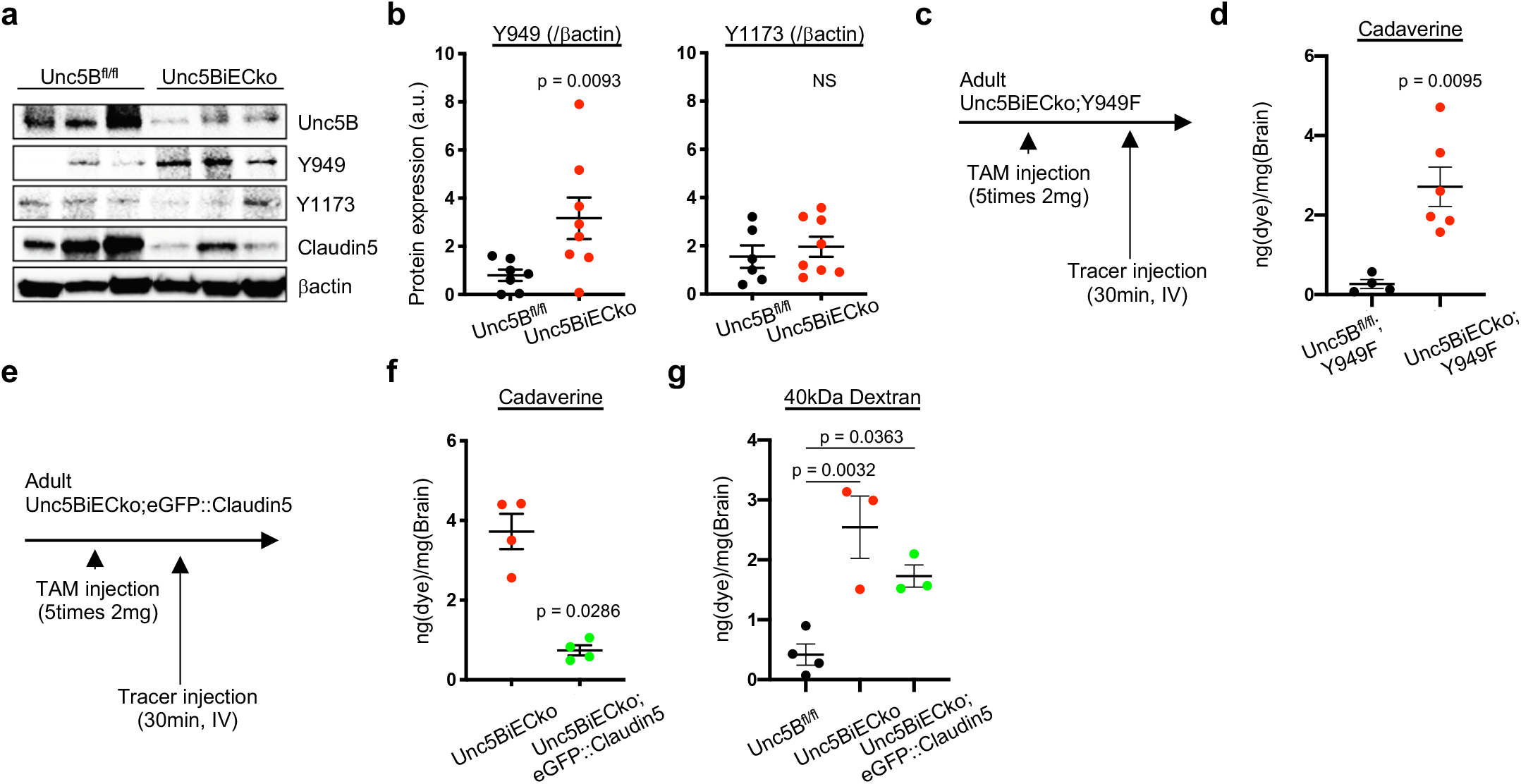
(a, b) Western blot and quantification of Vegfr2 signaling in adult *Unc5B^fl/fl^* and Unc5BiECko brain protein extracts, n > 7 mice per group. (c, d) Quantification of brain cadaverine content in adult mice, 7 days after TAM injection and 30 min after i.v. cadaverine injection (n > 4 mice per group). (e-g) Quantification of brain dye content in adult mice, 7 days after TAM injection and 30 min after i.v. cadaverine or 40kDa dextran injection (n > 3 mice per group). All data are shown as mean+/−SEM. NS: non-significant. Mann-Whitney U test was performed for statistical analysis between two groups. ANOVA followed by Bonferroni’s multiple comparisons test was performed for statistical analysis between multiple groups.

**Supplemental Figure 5:**
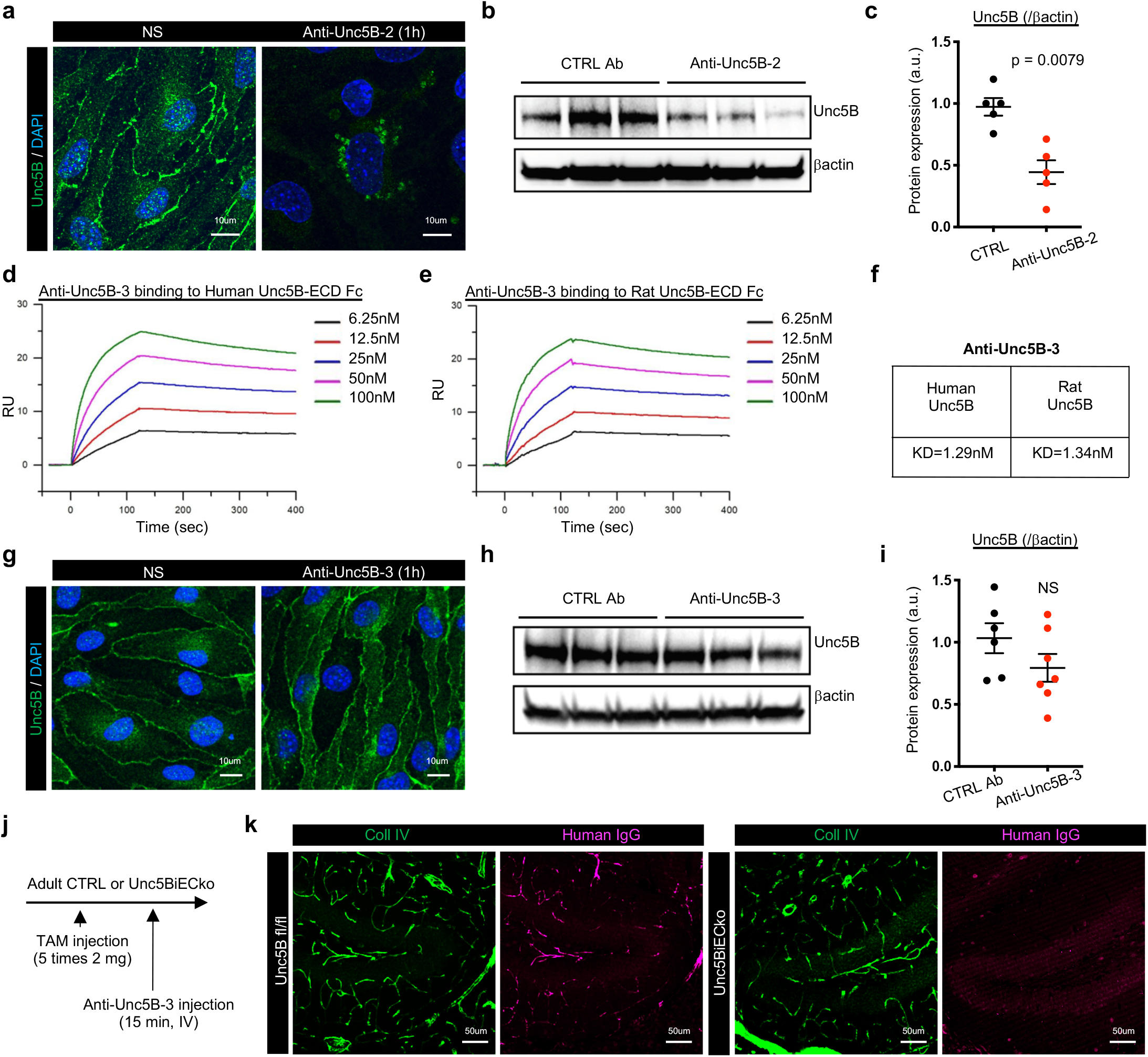
(a) Unc5B immunofluorescence detected with a commercial anti-Unc5B antibody and confocal imaging of confluent monolayers of mouse brain ECs (Bend3) treated or not with anti-Unc5B-2 for 1 h. (b,c) Western blot and quantification on brain protein extracts from mice i.v injected with CTRL or anti-Unc5B-2 antibodies (1 h, 10 mg/kg) (n = 5 mice per group). (d-f) Surface Plasmon Resonance measurements of anti-Unc5B-3 binding to human and rat Unc5B. (g) Unc5B immunofluorescence detected with a commercial anti-Unc5B antibody and confocal imaging of confluent monolayers of mouse brain ECs (Bend3) treated or not with anti-Unc5B-3 for 1 h. (h,i) Western-blot and quantification on brain protein extracts from mice i.v. injected with CTRL or anti-Unc5B-3 antibodies (1 h, 10 mg/kg) (n = 5 mice per group). (j,k) Anti-Unc5B-3 was i.v. injected in *Unc5B^fl/fl^* or Unc5BiECko mice for 20min, mice were perfused and anti-Unc5B-3 binding was revealed by immunofluorescence on brain vibratome section using an anti-human IgG antibody. All data are shown as mean+/−SEM. NS: non-significant, Mann-Whitney U test was performed for statistical analysis.

**Supplemental Figure 6:**
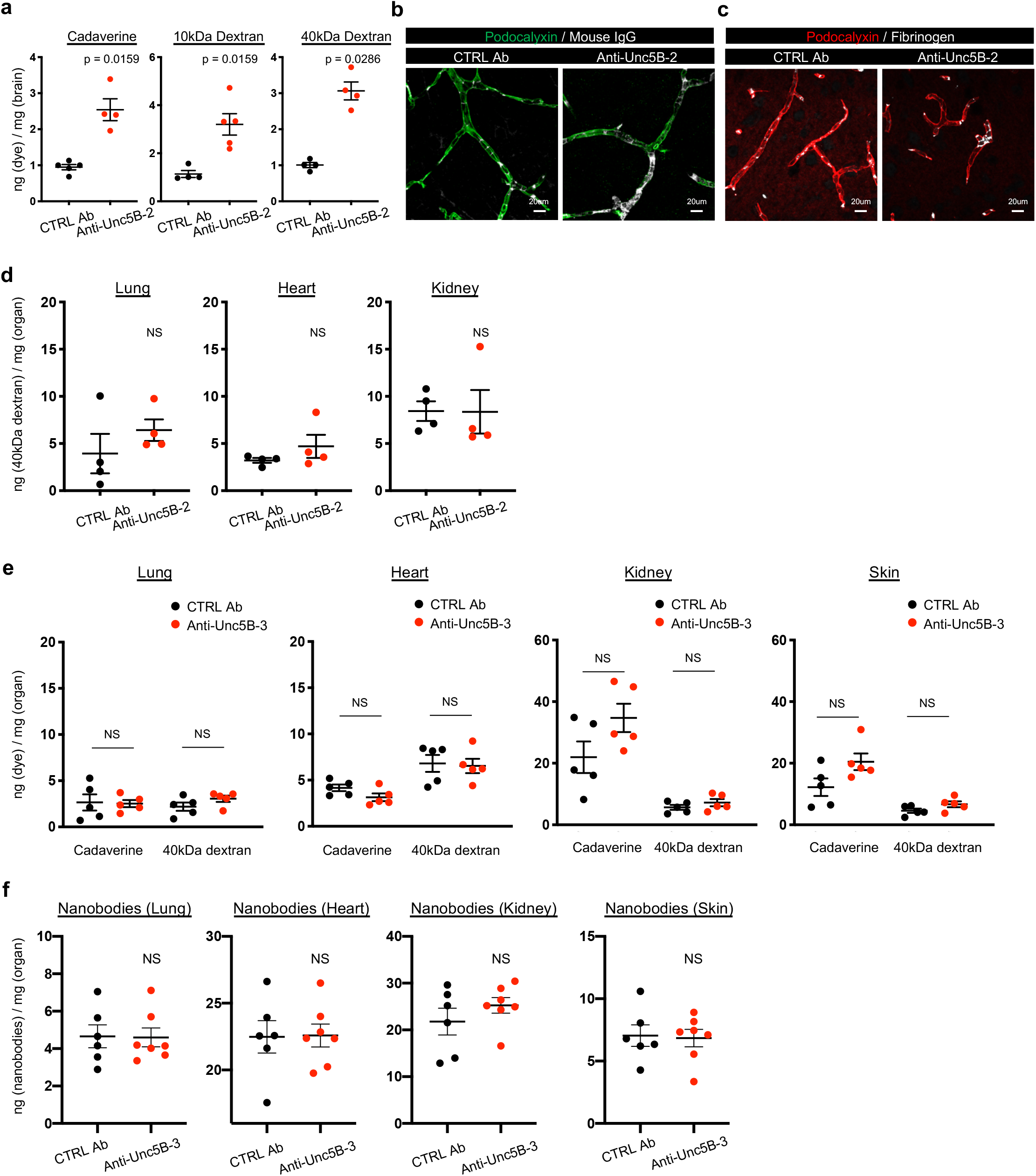
(a) Quantification of brain dye content 30min after injection of dyes with increasing MW and 1h after CTRL or anti-Unc5B-2 i.v. injection (10 mg/kg) (n > 4 mice per group). (b,c) Immunofluorescence of blood vessels (podocalyxin) and endogenous IgG and fibrinogen on brain vibratome sections 1 h after CTRL or anti-Unc5B-2 i.v. injection (10 mg/kg). (d,e) Quantification of organ dye content 1 h after i.v. CTRL or anti-Unc5B-2 or anti-Unc5B-3 injection (10 mg/kg) and 30min after i.v injection of dyes with increasing MW in adult mice (n > 4 mice per group). (f) Quantification of organ nanobody content 1 h after i.v CTRL or anti-Unc5B-3 injection (10 mg/kg) and 30 min after i.v nanobody injection (n = 5 mice per group). All data are shown as mean+/−SEM. NS: non-significant, Mann-Whitney U test was performed for statistical analysis.

**Supplemental Figure 7:**
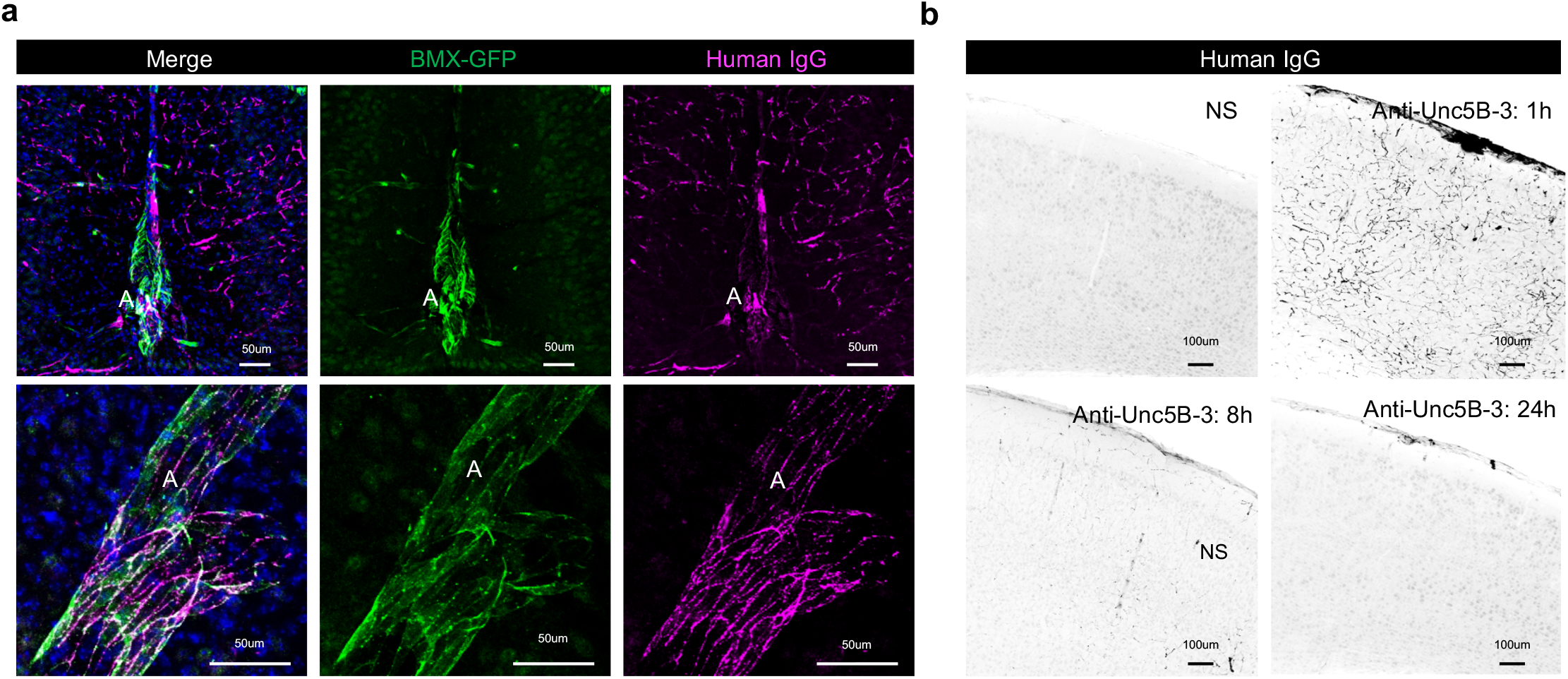
(a) TAM was injected for 5 days in *BMXCre^ERT2^-mTmG* mice followed by anti-Unc5B-3 i.v. injection for 15 min. After cardiac perfusion, anti-Unc5B binding was revealed by immunofluorescence using anti-human IgG antibody followed by confocal imaging. (b) Immunofluorescence staining of anti-human IgG antibody on adult brain vibratome section from mice i.v. injected with anti-Unc5B-3 (10mg/kg) for 1, 8 or 24 h.

**Supplemental Figure 8:**
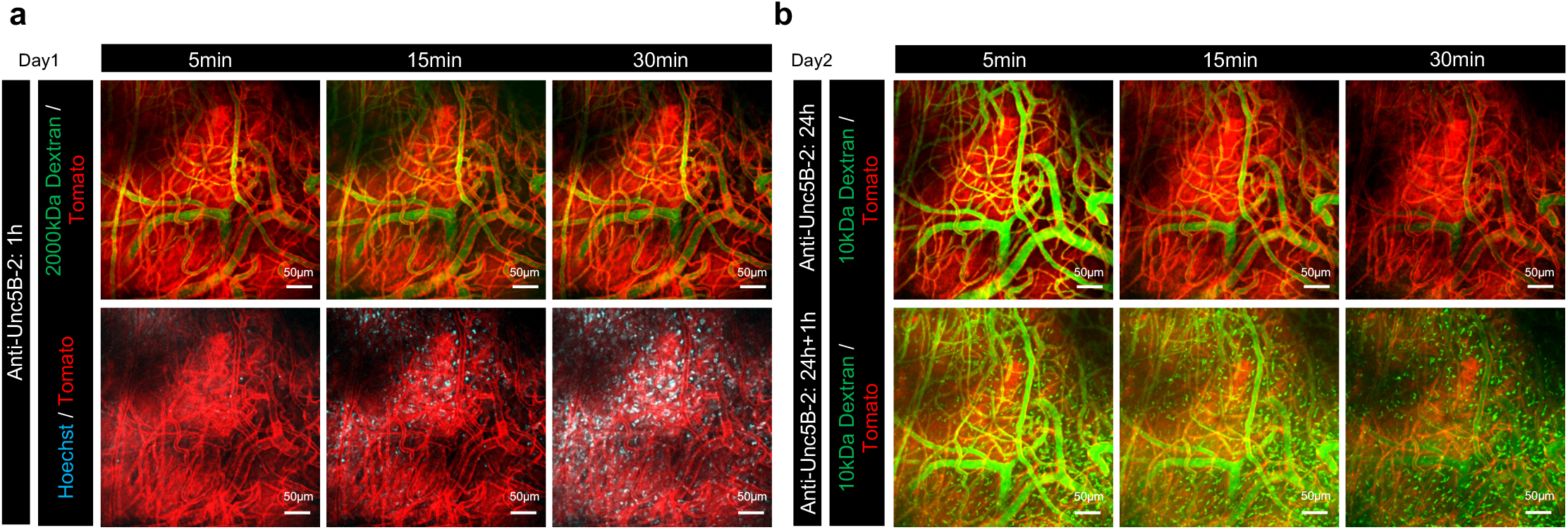
(a) Two-photon live imaging of *C57BL/6; ROSAmTmG* mice 1h after i.v. injection of anti-Unc5B-2 (1 h, 10 mg/kg) and 5, 15 or 30 min after i.v. injection of 2000kDa FITC-dextran and 560Da Hoechst. Note leakage of Hoechst 15 to 30min after i.v. injection while 2000kDa FITC-dextran outlined the brain vasculature. (b) Two-photon live imaging of mice that were treated with the anti-Unc5B-2 antibody 24 h earlier, 5, 15 and 30 min after i.v. injection of 10kDa FITC-dextran. Note the absence of BBB leakage. Next, mice received a second i.v. injection of anti-Unc5B-2 (10 mg/kg) for 1h followed by another i.v. injection of 10kDa FITC-dextran and two-photon live imaging revealed BBB leakage.

**Supplemental Figure 9:**
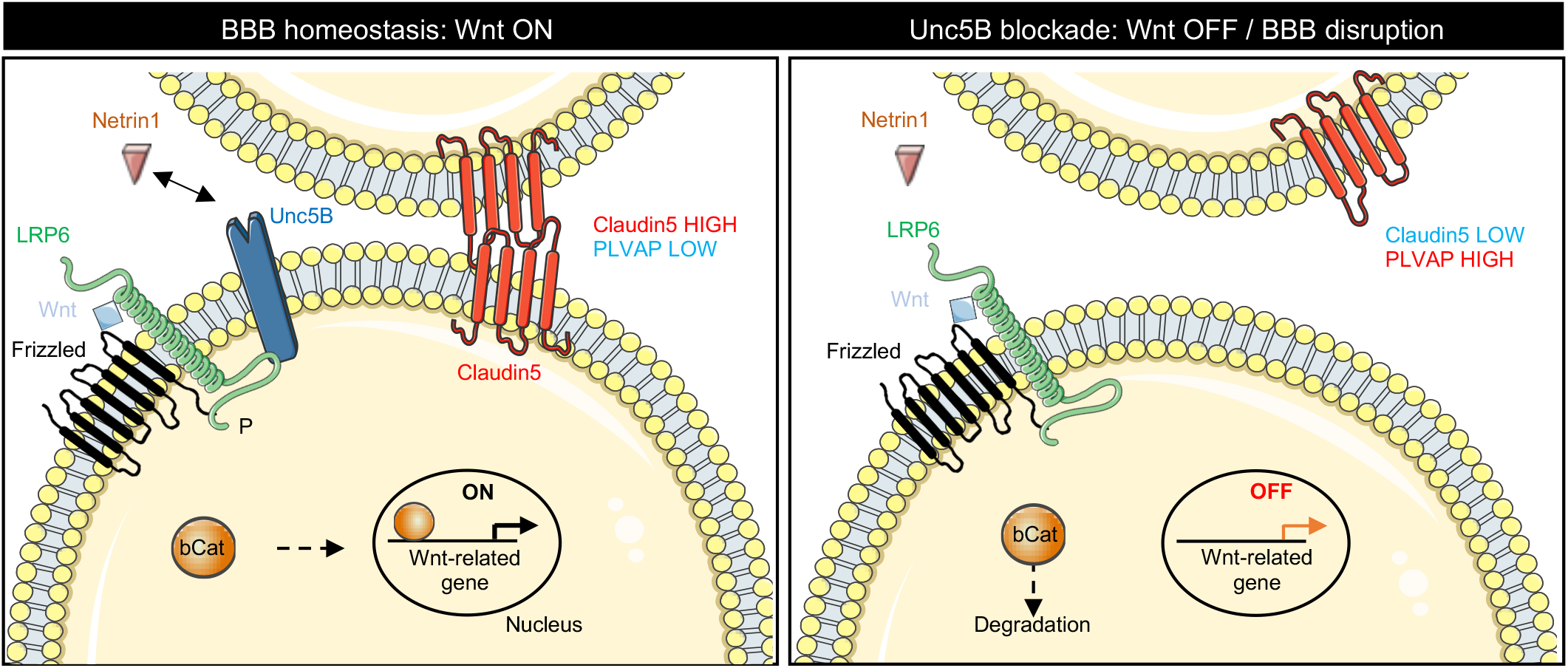
Netrin1 binding to endothelial Unc5B regulates Wnt/β-catenin signaling and BBB integrity. In the absence of Netrin1-Unc5B signaling the Wnt/β-catenin signaling is disrupted which induced loss of Claudin5 along with increased PLVAP expression and BBB leakage.

